# WgLink: reconstructing whole-genome viral haplotypes using *L*_0_ + *L*_1_-regularization

**DOI:** 10.1101/2020.08.14.251835

**Authors:** Chen Cao, Matthew Greenberg, Quan Long

## Abstract

Many tools can reconstruct viral sequences based on next generation sequencing reads. Although existing tools effectively recover local regions, their accuracy suffers when reconstructing the whole viral genomes (strains). Moreover, they consume significant memory when the sequencing coverage is high or when the genome size is large. We present WgLink to meet this challenge. WgLink takes local reconstructions produced by other tools as input and patches the resulting segments together into coherent whole-genome strains. We accomplish this using an *L*_0_ + *L*_1_-regularized regression synthesizing variant allele frequency data with physical linkage between multiple variants spanning multiple regions simultaneously. WgLink achieves higher accuracy than existing tools both on simulated and real data sets while using significantly less memory (RAM) and fewer CPU hours. Source code and binaries are freely available at https://github.com/theLongLab/wglink.

## INTRODUCTION

In the viral sequencing projects based on next-generation sequencing (NGS) instruments, researchers typically sample a collection of viruses in tissue rather than targeting individual viruses. As a result, sequencing reads represent a mixture of viral particles reflecting the within-host viral population, the constituents of which are called viral quasispecies. Reconstructing haplotypes from this mixture is an important but difficult computational problem.

Haplotype reconstruction from pooled NGS data is a prerequisite for many downstream biological analyses focusing on individual strains. This has led bioinformaticians to develop tools facilitating such reconstruction. These tools employ an range of computational and statistical techniques, e.g. graph algorithms (1, 2) regularization (3), Tenser factorization (4), Dirichlet processes (5), and Bayesian methods (6).

Many tools can effectively reconstruct haplotypes locally at the gene level (4, 6). However, the accuracy of global reconstruction, i.e. at the level of the whole viral genome (strain) remains low. Moreover, the techniques for such global reconstruction consume significant memory resources. There is much room for improvement.

In this work, we present WgLink, a novel tool using regularized regression to stitch together regional haplotypes into global ones. As input, WgLink takes regional haplotypes produced by existing tools, the reference genome of the focal species, and the sequencing reads of a sample. It outputs whole viral genome haplotypes present in the sample together with their within-host frequencies. WgLink’s novelty lies both in its design, using a regression incorporating both allele frequencies and physical linkage disequilibrium (PLD) observed in sequencing reads, and its implementation, solving this regression problem using a state-of-the-art tool for extra-fast *L*_0_-regularization (7) (**Methods**). As a result, WgLink generates more accurate viral haplotypes using less memory and fewer CPU hours.

## METHOD

### The WgLink algorithm

Based on regional haplotypes generated by some other tool, WgLink first generates all possible whole-genome haplotypes using a breadth-first search (**Supplementary Notes**). To collect allele frequency and LD information in a consistent file format, WgLink uses BWA (8) to map reads back to the reference genome. Subsequently, WgLink performs a regularized regression to estimate the haplotype frequencies of all candidates.

As indicated above, the design of this regularized regression contains **two key innovations**. First is the synthesis of allele frequencies at individual sites and the LD between pairs of sites in the context of a single regression problem. We explain why these innovations lead to the claimed improvements. Many existing tools aiming to reconstruct haplotypes targeting different species use allele frequencies only (3, 9, 10). More specifically: Letting Y denote the vector of within-host frequencies of all genetic variants and *X* = (*X*_1_, *X*, …, *X*_*k*_) denote the potential haplotypes, represented by vectors of 0s and 1s (representing reference and alternate alleles), then the linear regression of *Y*~ ∑ *β*_*i*_*X*_*i*_ gives an estimate of the haplotype frequencies *β*_1_, *β*_2_, …, *β*_*k*_. (Supplementary Fig. S1). Note that in a regularized form, most of the *β*_*i*_ are 0. The sample size for solving this regression is the number of genetic variants. This model is useful only if there are sufficient polymorphic sites to distinguish the various haplotypes. However, this linear regression approach using allele frequency alone is insufficient for distinguishing haplotypes that differ subtly, e.g. by a few recombination events. For instance, the actual haplotypes and frequencies in Fig. 1, (x1:1/6, x2:2/6, x3:3/6), will have the same allele frequencies as the configuration (x4:1/6, x5:2/6, x6:3/6) and, thus, won’t be distinguishable by regression on allele frequency only. As remedy for this insensitivity, WgLink incorporates a quantity derived from pairs of alleles on the same sequencing read (or paired-end reads) that we call Physical Linkage Disequilibrium, or PLD (in contrast to statistical LD in population genetics). More precisely, for any two sites, one sample will be added as follows: the proportion of reads carrying allele 1 on both sites (out of all reads covering both sites) will be added as a Y-value. The corresponding xi value will be set to 1 if the corresponding i-th potential haplotype contains allele 1 on both sites and to 0, otherwise. As such, samples capturing the PLD, i.e., the blue diamonds in Fig. 1, calculated in the left panel and used as samples in the right panel (line 5-10), will be added to the regression. For highly polymorphic viruses such as HIV, the sequencing reads contain a large number of PLDs -- about 50-100 times of number of genetic variants -- leading to a better estimate of the number and frequencies of haplotypes.

**Fig. 1:**
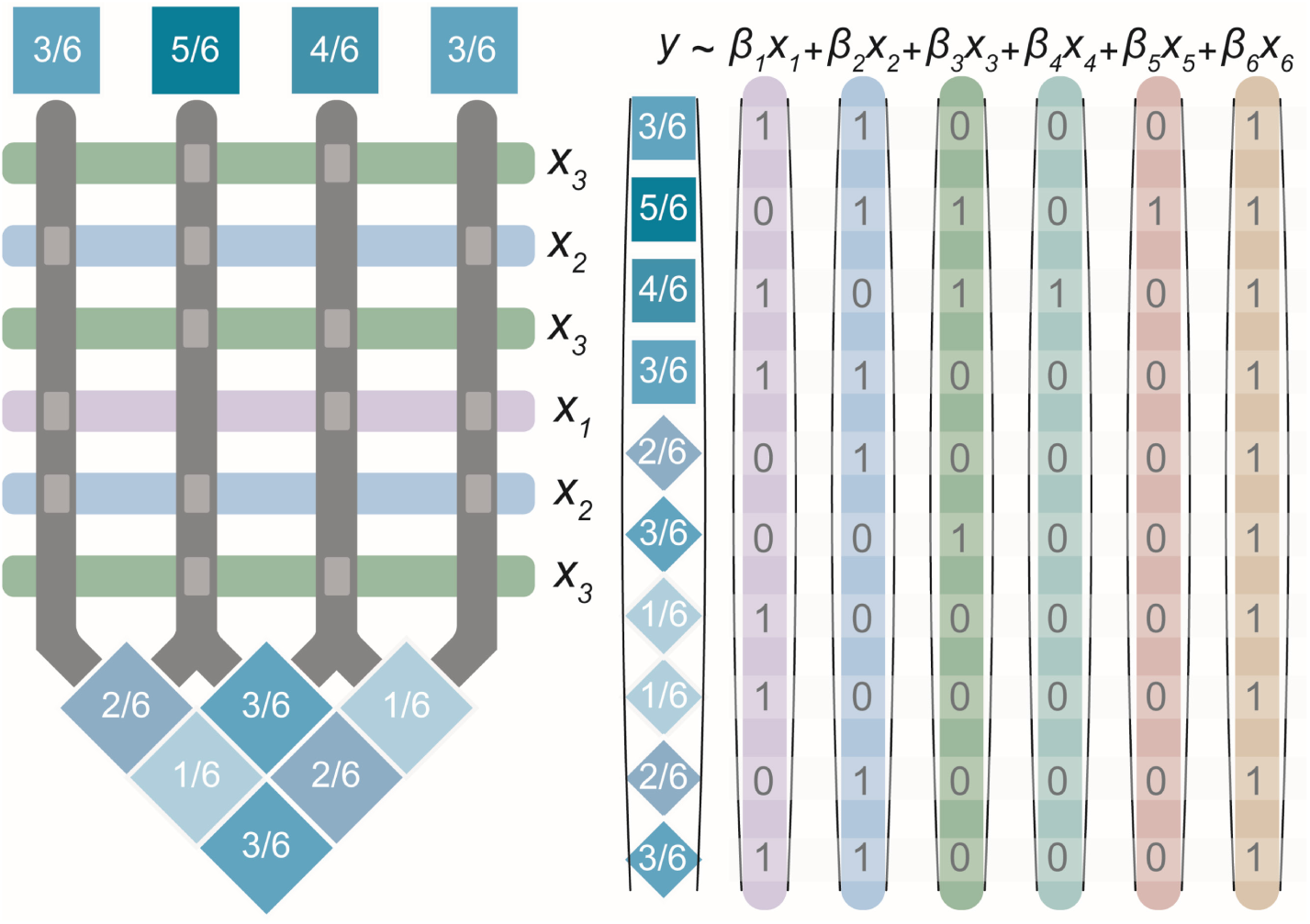
Regression integrating both allele frequency and physical LD. Colored bars on the left represents reads belonging to three actual haplotypes *x*_1_, *x*_2_, and *x*_3_, corresponding to the vectors on the right. The vectors *x*_4_, *x*_5_, and *x*_6_ represent other potential haplotypes with actual frequencies 0. Blue squares denote allele frequencies and diamonds denote physical LDs.

The second innovation is adding of a mixture of *L*_0_ and *L*_1_ regularization to the regression problem. Existing tools use *L*_1_ regularization (i.e., LASSO) only (9), which does not discriminate feasible solutions for which the haplotype frequencies sum (correctly) to 1. For example, an *L*_1_-regularized regression does not favor a correct solution of 2 haplotypes, each with frequency 0.5, over a competing, incorrect solution of 10 haplotypes, all with frequency being 0.1, because of the penalty term being 1.0 in both cases. RegressHaplo attempted to address this using a revised *L*_1_-regularization term. To fundamentally solve this problem, we add an *L*_0_-regularization term, imposing a much stiffer penalty on high-complexity models (**Supplementary Notes**). *L*_0_-regularized regression is a non-convex optimization problem whose solution was, until recently, extremely slow. This changed with the release of L0Learn package (7), an exciting breakthrough in the field, that can solve *L*_0_-regularized regression problems efficiently. Although the L0Learn paper does not present a precise estimate of time-complexity, our benchmarking shows that our L0Learn-based implementation of WgLink is significantly faster than other tools (Supplementary Table S1).

### Comparison with other tools

We found that WgLink worked the best with regional haplotype inputs constructed by TenSQR (4) and aBayesQR (6). So in the comparisons, WgLink is supported by one of these two tools. We compared the accuracy of WgLink against four popular tools: TenSQR, aBayesQR, RegressHaplo (3) and CliqueSNV(1). Specifically, we compare the results of directly running the other tools on the whole genome with those of WgLink computed by generating local haplotypes (using TenSQR or aBayesQR) in tiling windows and then linking them into the whole genome. To thoroughly benchmark the alternative tools, we assessed the performance using four criteria: (1) the difference between gold-standard allele frequencies and reconstructed frequencies assessed by Jensen–Shannon divergence (JSD); (2) the number of differences between gold-standard haplotypes and reconstructed haplotypes; (3) the deviation of the total number of reconstructed haplotypes from the actual number of haplotypes. The comparisons were conducted on both simulated and real data sets with known haplotype composition(11). In the simulations, we tested the combinations of low/high coverages (2.5K or 10K), low/high sequencing errors (0.1% or 0.4%), and long/short read-length (150bp or 250bp). Details of simulations, comparison criteria, and application of alternative tools are presented in **Supplementary Notes**.

## RESULT

For accuracy, WgLink, integrated with either TenSQR or aBayesQR, outperforms alternative tools with a large margin in terms of all three of the above criteria. This observation persists regardless of sequencing coverage, sequencing error, or read-length (Supplementary Figures S2 - S9). The same trend is observed in real data, for which WgLink (with either TenSQR or aBayesQR) achieved the correct number of haplotypes, very high reconstruction rate (>93% on all polymorphic sites), and very close frequencies (JSD <0.1), much better than the other tools that were able to finish (Supplementary Table S2). Moreover, WgLink used significantly less memory and CPU time (Supplementary Table S1) in our HPC cluster (specification in **Supplementary Notes**).

## SUPPLEMENTARY MATERIALS

### Detailed algorithm for the breadth-first search

The local haplotypes produced by the other reconstruction tool are linked into potential global haplotypes using a breadth-first search (BFS) algorithm. As haplotypes are inferred for each tiling window, haplotypes from adjacent windows will overlap over about half of their sequence length. If two overlapping haplotypes match on this overlap, they are stitched together to form a longer haplotype. Since the overlapping regions may not be identical due to errors in the previous steps, two haplotypes may be stitched together as long as the number of mismatches in their overlapping region is less than a user-specified cutoff parameter.

Next, global haplotypes are determined by traversing a tree structure whose nodes are the locally stitched haplotypes. The *i*-th level of the tree represents the *i*-th genomic region. We draw an edge between two nodes in neighboring levels if and only if their corresponding haplotypes can be stitched together.

Paths that traverse the tree enumerated using BFS, represent candidate global haplotypes, as stitched together local ones. These are vetted further in WgLink’s regularized regression phase.

### Real Data, Simulations, and HPC specs

#### Real data

For benchmarking on real data, we used a public dataset containing a five-strain(1). Specifically, this dataset describes an in vitro generated quasispecies population consisting of 5 known HIV-1 strains (HIV-1_HXB2_, HIV-1_89:6_, HIV-1_JR-CSF_, HIV-1_NL4-3_ and HIV-1_YU2_) with pairwise distances between 2.61-8.45%; the estimated frequency for the five strains are 22.6% (HIV-1_89:6_), 10.0% (HIV-1_HXB2_), 29.6% (HIV-1_JR-CSF_), 26.9% (HIV-1_NL4-3_) and 10.9% (HIV-1_YU2_).

#### Simulations

We used the above five-strain as templates for simulated data. In real world, haplotype abundance is unknown and random. To reflect this, we randomly generated gold-standard frequencies for each haplotype. Based on the gold-standard haplotypes (i.e., the five strains in real data) and their frequencies, DWGSIM (2) (https://github.com/nh13/DWGSIM) was used to generate paired-end reads for a mixture of the haplotypes. We varied three factors in our construction: sequencing error (low = 0.1%, high = 0.4%), read length (low = 150bp, high = 250bp) and average sequencing coverage per base pair (low = 2500X, high = 10000X), giving a total eight combinations (see Supplementary Figures 2 – 9). For each combination of simulation parameters, we performed fifty (50) simulation rounds and plotted the averages. To avoid unusually low values, we discarded any haplotype with abundance less than 1%, replacing it with a new simulation.

#### Data processing

For both real and simulated data, we used BWA (version 0.7.15) (3) and SAMTools (4) (version 1.4) mapping and sorting reads, respectively. We used HXB2, the standard reference genome for HIV studies, as the reference genome.

##### Specification of the high-performance computing (HPC) cluster nodes

CPU: 2 sockets, each with 14 cores with Intel(R) Xeon(R) CPU E5-2690 v4 @ 2.60GHz Memory: 188 GB DDR4

Network: 40Gb/s infiniband

Storage: 1.3PB GPFS storage shared by all nodes.

### Parameters and commands for different tools

We used the default parameters of TenSQR, aBayesQR is WgLink. According to the TenSQR(5) and aBayesQR manual(6), specified min_mapping_qual, min_read_length and MEC improvement threshold were used for the real data analysis.

### WgLink_TenSQR parameters and explanation

~~~
*#The name of the project.*
Proj_Name = Project_Name
*#Absolute paths for fastq files.*
Fastq_1_Path = /Path/to/Test.1.fastq
Fastq_2_Path = /Path/to/Test.2.fastq
*#Absolute path for reference file.*
Reference_Seq = /Path/to/reference.fa
*#File locations: output files directory.*
Output_Path = /Path/to/Output_Folder
*#Reconstruction Start Position, 2*read length*
Reconstruction_Start = 501
*#Reconstruction End Position, reference length-2*read length*
Reconstruction_End = 9219
*#The length of each region for divide and conquer*
Region_Length = 500
*#Mapping quality cutoff*
Min_Mapping_Qual = 40
*#Read length cutoff*
Min_Read_Length = 100
*#Maximum insert read length*
Max_Insert_Length = 1000
*#Estimated sequencing error*
Sequence_Err = 0.1
*#SNV_Cutoff, please refer to TenSQR user manual*
SNV_Cutoff = 0.01
*#MEC_Improvement_Cutoff, please refer to TenSQR user manual*
MEC_Improvement_Cutoff = 0.0312
*#Initial_Population_Size, please refer to TenSQR user manual*
Initial_Population_Size = 5
*#The weight of the constraint Sigma freq_i = 1, where freq_i is in the in-pool frequency for haplotype_i.*
Regression_One_Vector_Weight = 50.0
*#The weight of the constraints Sigma freq_i * h_ij = MAF_j (j is the SNP index and i is the haplotype index)*
Regression_Hap_MAF_Weight = 5.0
*#The weight for PLD (specifically, the probability for both SNP_k and SNP_j being the alternate allele)*
Regression_Hap_LD_Weight = 1.0
*#Maximum SNP mismatch ratio tolerance in region extextion*
BFS_Mismatch_Tolerance_Rate = 0.1
Number_Threads = 8
*#Number of maximum selected haplotypes to generate higher level potential haplotypes for following L0L1 regression.*
Maximum_Haps_R = 20
*#The minimum regularization gamma penalty for L0L1 regression.*
Regression_Gamma_Min =0.0001
*#The maximum regularization gamma penalty for L0L1 regression.*
Regression_Gamma_Max =0.1
*#The number of gamma values beween Regression_Gamma_Min and Regression_Gamma_Max for L0L1 regression.*
Regression_n_Gamma = 10
*#The lambda penalty for L0L1 regression.*
Regression_Lambda = 0.002
*#The minimum frequency cutoff for haplotype.*
Min_Hap_Freq = 0.01
*#Number of maximum potential haplotypes for L0L1 regression.*
Max_L0L1_Regional_Haps = 1000
*#Absolute paths for bwa.*
bwa = /Path/to/bwa
*#Absolute paths for Rscript.*
Rscript_path = /Path/to/Rscript
*#Absolute paths for ExtractMatrix.*
ExtractMatrix = /Path/to/ExtractMatrix
*#Absolute paths for TenSQR.py.*
TenSQR = /Path/to/TenSQR.py
*#Absolute paths for PYTHON3.*
PYTHON = /Path/to/PYTHON3
~~~

### WgLink_aBayesQR parameters and explanation

~~~
*#The name of the project.*
Proj_Name = Project_Name
*#Absolute paths for fastq files.*
Fastq_1_Path = /Path/to/Test.1.fastq
Fastq_2_Path = /Path/to/Test.2.fastq
*#Absolute path for reference file.*
Reference_Seq = /Path/to/reference.fa
*#File locations: output files directory.*
Output_Path = /Path/to/Output_Folder
*#Reconstruction Start Position, 2*read length*
Reconstruction_Start = 501
*#Reconstruction End Position, reference length-2*read length*
Reconstruction_End = 9219
*#The length of each region for divide and conquer*
Region_Length = 500
*#Mapping quality cutoff*
Min_Mapping_Qual = 40
*#Read length cutoff*
Min_Read_Length = 100
*#Maximum insert read length*
Max_Insert_Length = 1000
*#Estimated sequencing error*
Sequence_Err = 0.1
*#SNV_Cutoff, please refer to aBayesQR user manual*
SNV_Cutoff = 0.01
*#MEC_Improvement_Cutoff, please refer to aBayesQR user manual*
MEC_Improvement_Cutoff = 0.0312
*#The weight of the constraint Sigma freq_i = 1, where freq_i is in the in-pool frequency for haplotype_i.*
Regression_One_Vector_Weight = 50.0
*#The weight of the constraints Sigma freq_i * h_ij = MAF_j (j is the SNP index and i is the haplotype index)*
Regression_Hap_MAF_Weight = 5.0
*#The weight for PLD (specifically, the probability for both SNP_k and SNP_j being the alternate allele)*
Regression_Hap_LD_Weight = 1.0
*#Maximum SNP mismatch ratio tolerance in region extextion*
BFS_Mismatch_Tolerance_Rate = 0.1
Number_Threads = 8
*#Number of maximum selected haplotypes to generate higher level potential haplotypes for following L0L1 regression.*
Maximum_Haps_R = 20
*#The minimum regularization gamma penalty for L0L1 regression.*
Regression_Gamma_Min =0.0001
*#The maximum regularization gamma penalty for L0L1 regression.*
Regression_Gamma_Max =0.1
*#The number of gamma values beween Regression_Gamma_Min and Regression_Gamma_Max for L0L1 regression.*
Regression_n_Gamma = 10
*#The lambda penalty for L0L1 regression.*
Regression_Lambda = 0.002
*#The minimum frequency cutoff for haplotype.*
Min_Hap_Freq = 0.01
*#Number of maximum potential haplotypes for L0L1 regression.*
Max_L0L1_Regional_Haps = 1000
*#Absolute paths for bwa.*
bwa = /Path/to/bwa
*#Absolute paths for Rscript.*
Rscript_path = /Path/to/Rscript
*#Absolute paths for aBayesQR.*
aBayesQR = /Path/to/aBayesQR
~~~

### TenSQ parameters

~~~
filename of reference sequence (FASTA): /Path/to/HIV_HXB2.fa
filname of the aligned reads (sam format): /Path/to/Project_Name.sam
SNV_thres: 0.01
*#reconstruction_start= 501 when read length =250bp*
reconstruction_start: 301
*#reconstruction_stop= 9219 when read length =250bp*
reconstruction_stop: 9419
*#min_mapping_qual = 60 when read length =250bp*
min_mapping_qual: 40
*#min_read_length = 150 when read length =250bp*
min_read_length: 100
max_insert_length: 1000
characteristic zone name: Project_Name
seq_err (assumed sequencing error rate(%)): 0.1
MEC improvement threshold: 0.0312
initial population size: 5
~~~

### aBayesQR parameters

~~~
filename of reference sequence (FASTA): /Path/to/HIV_HXB2.fa
filname of the aligned reads (sam format): /Path/to/Project_Name.sam
paired-end (1 = true, 0 = false): 1
SNV_thres: 0.01
*#reconstruction_start= 501 when read length =250bp*
reconstruction_start: 301
*#reconstruction_stop = 9219 when read length =250bp*
reconstruction_stop: 9419
*#min_mapping_qual = 60 when read length =250bp*
min_mapping_qual: 40
*#min_read_length = 150 when read length =250bp*
min_read_length: 100
max_insert_length: 1000
characteristic zone name: Project_Name
seq_err (assumed sequencing error rate(%)): 0.1
MEC improvement threshold: 0.0312
~~~

### RegressHaplo R command

~~~
library(“RegressHaplo”)
*#read length= 150bp*
full_pipeline(“Project_Name/Test.bam”, “.”, start_pos=301, end_pos=9419, num_trials=701)
*#read length= 250bp*
full_pipeline(“Project_Name/Test.bam”, “.”, start_pos=501, end_pos=9219, num_trials=701)
~~~

### CliqueSNV command

~~~
/Path/to/java -Xmx80G -jar /Path/to/clique-snv.jar -m snv-illumina -tf 0.01 -in Project_Name.bam -threads 8 -log
~~~

### Comparison criteria

#### Estimated number of haplotypes

The estimated numbers of haplotypes are the numbers of predicted haplotypes by different tools (including WgLink and alternative tools). The error in an estimate is simply the difference between the prediction and the actual number.

#### Reconstruction Rate (two versions)

Reconstruction rate is, by definition, the number of mismatches between the gold-standard halophytes and the reconstructed ones. Since these may differ in number, we need an explicit mapping between gold-standard and reconstructed haplotypes to facilitate the comparison. There are two natural ways constructing such: mapping the gold-standard haplotypes to reconstructed haplotypes or mapping the reconstructed haplotypes to gold-standard haplotypes, corresponding to Reconstruction Rates 1 and 2 (see below) respectively.

##### Reconstruction Rate 1

is defined as

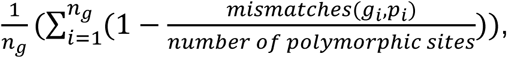

where the *g*_*i*_, *i* = 1, …, *n*_*g*_, are the gold-standard haplotypes, *p*_*i*_ is the predicted haplotype closest to *g*_*i*_ in terms of minimum Hamming distance, and *mismatches* is the number of mismatch sites between *g*_*i*_ *and p*_*i*_.

##### Reconstruction Rate 2

Reconstruction Rate 2 is defined as

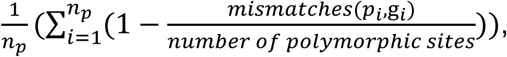

where the *p*_*i*_, *i* = 1, …, *n*_*p*_, are the predicted haplotypes, *g*_*i*_ is the gold-standard haplotype closest to *p*_*i*_ in terms of minimum Hamming distance, and *mismatches* is the number of mismatch sites between *g*_*i*_ *and p*_*i*_.

### Jensen–Shannon divergence (JSD)

The Jensen–Shannon divergence (JSD) evaluates the divergence between the frequencies of the simulated and reconstructed haplotype distributions and represents our tool’s ability to predict haplotype abundance. The probability distributions A and P represent the frequency distributions gold-standard and predicted haplotypes, respectively. JSD is defined as

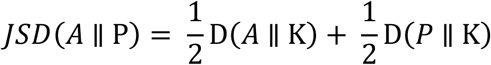

where

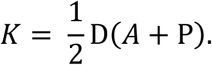

The quantity *D*(*A*||*P*) is the Kullback–Leibler divergence from A to P, defined by

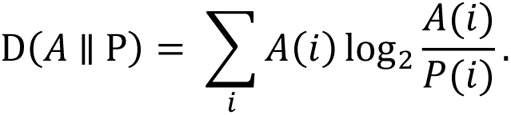

JSD is symmetric in A and P and takes values in the interval [0,1], the value 0 representing a perfect prediction.

---

**Supplementary Fig. 1.**
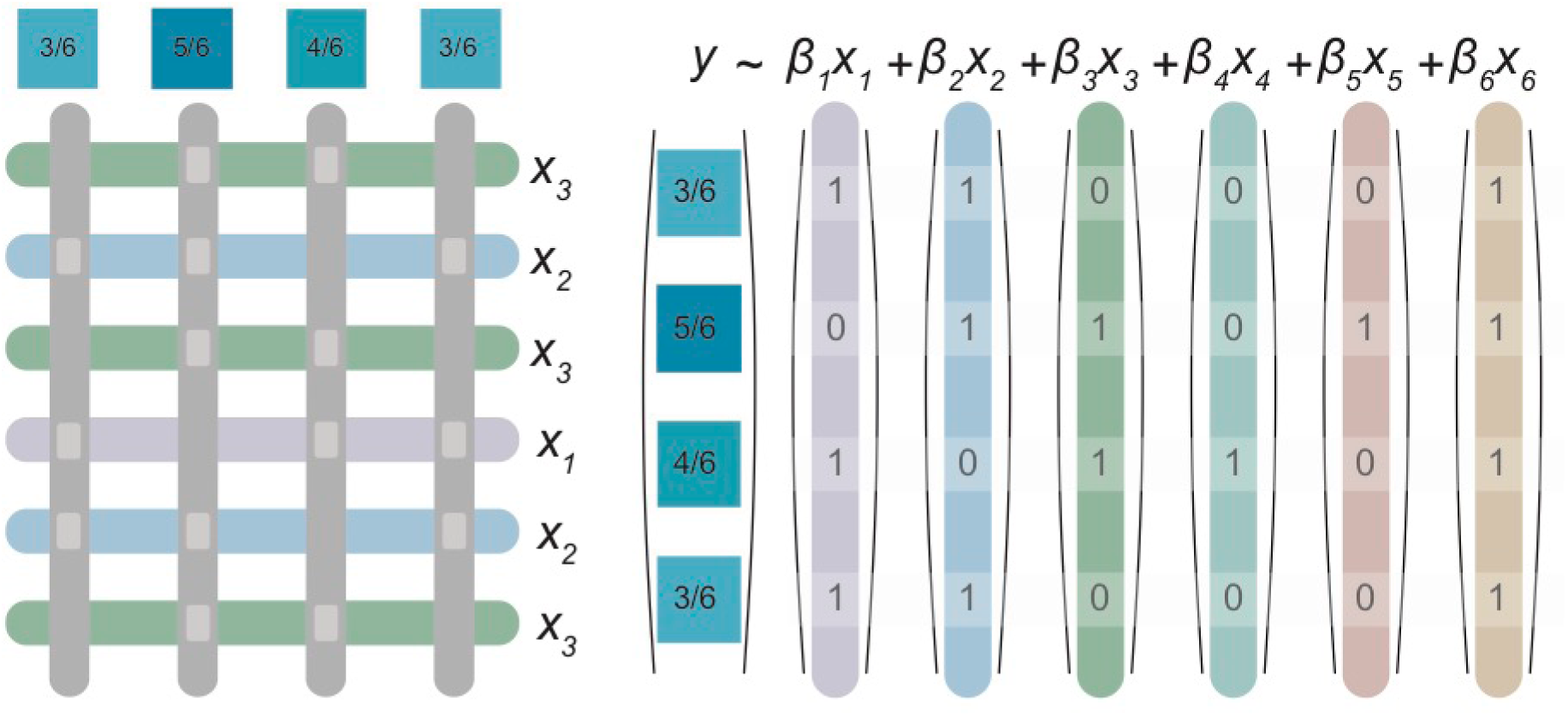
A naïve regression based on allele frequencies only. Colored bars on the left represents reads belongs to three actual haplotypes (x_1_, x_2_, and x_3_), corresponding to the vectors on the right. x_4_, x_5_, and x_6_ are potential haplotypes with actual frequencies 0. (As stated in the main text, without incorporating physical linkage disequilibrium, it can’t distinguish many incorrect configurations.)

**Supplementary Fig. 2:**
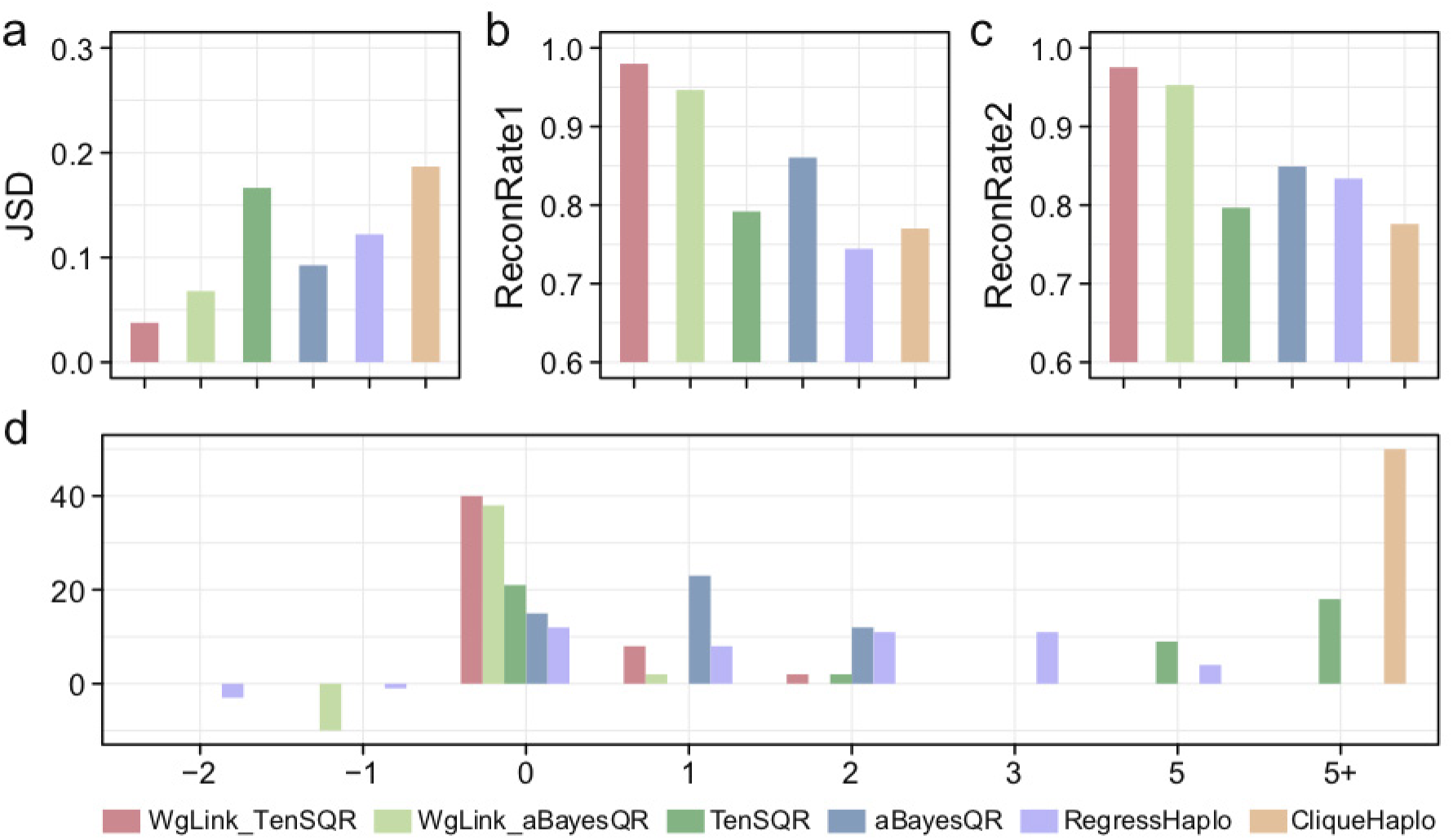
Comparison between different viral haplotype reconstruction tools based on the parameterization: 150bp read length, 2500X coverage, 0.1% sequencing error rate. Fifty rounds of simulations were conducted, and the averages were plotted. Panel **a** shows the accuracy for haplotype frequency (JSD) between truth and predicted (the smaller the better); Panels **b** and **c** show the accuracy for haplotype reconstruction, ReconRate1 and ReconRate2 based on two strategies of mapping the identities described in “Comparison criteria” (the larger the better). Penal **d** displaces the distribution of differences between the actual number of haplotypes and the predicted number. 0 means identical the predicted is identical to the truth (i.e., no error); and 5+ means the tools predicted 5 more haplotypes than truth (indicating large error).

**Supplementary Fig. 3:**
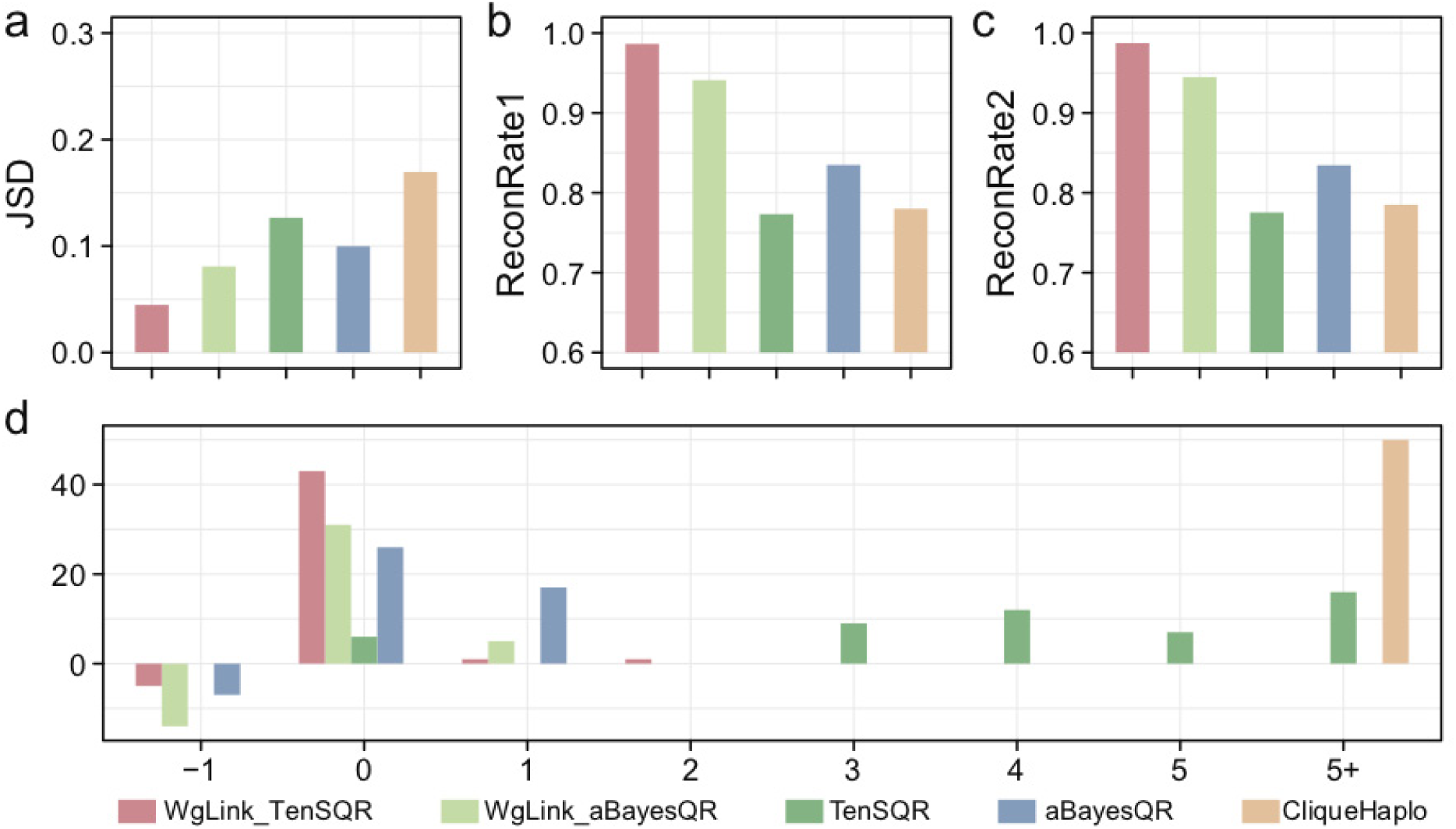
Comparison between different viral haplotype reconstruction tools based on the parameterization: 150bp read length, 2500X coverage, 0.4% sequencing error rate. Fifty rounds of simulations were conducted, and the averages were plotted. Panel **a** shows the accuracy for haplotype frequency (JSD) between truth and predicted (the smaller the better); Panels **b** and **c** show the accuracy for haplotype reconstruction, ReconRate1 and ReconRate2 based on two strategies of mapping the identities described in “Comparison criteria” (the larger the better). Penal **d** displaces the distribution of differences between the actual number of haplotypes and the predicted number. 0 means identical the predicted is identical to the truth (i.e., no error); and 5+ means the tools predicted 5 more haplotypes than truth (indicating large error). Only three alternative tools are displaced because RegressHaplo didn’t generate outcome when sequencing error rats is 0.4%.

**Supplementary Fig. 4:**
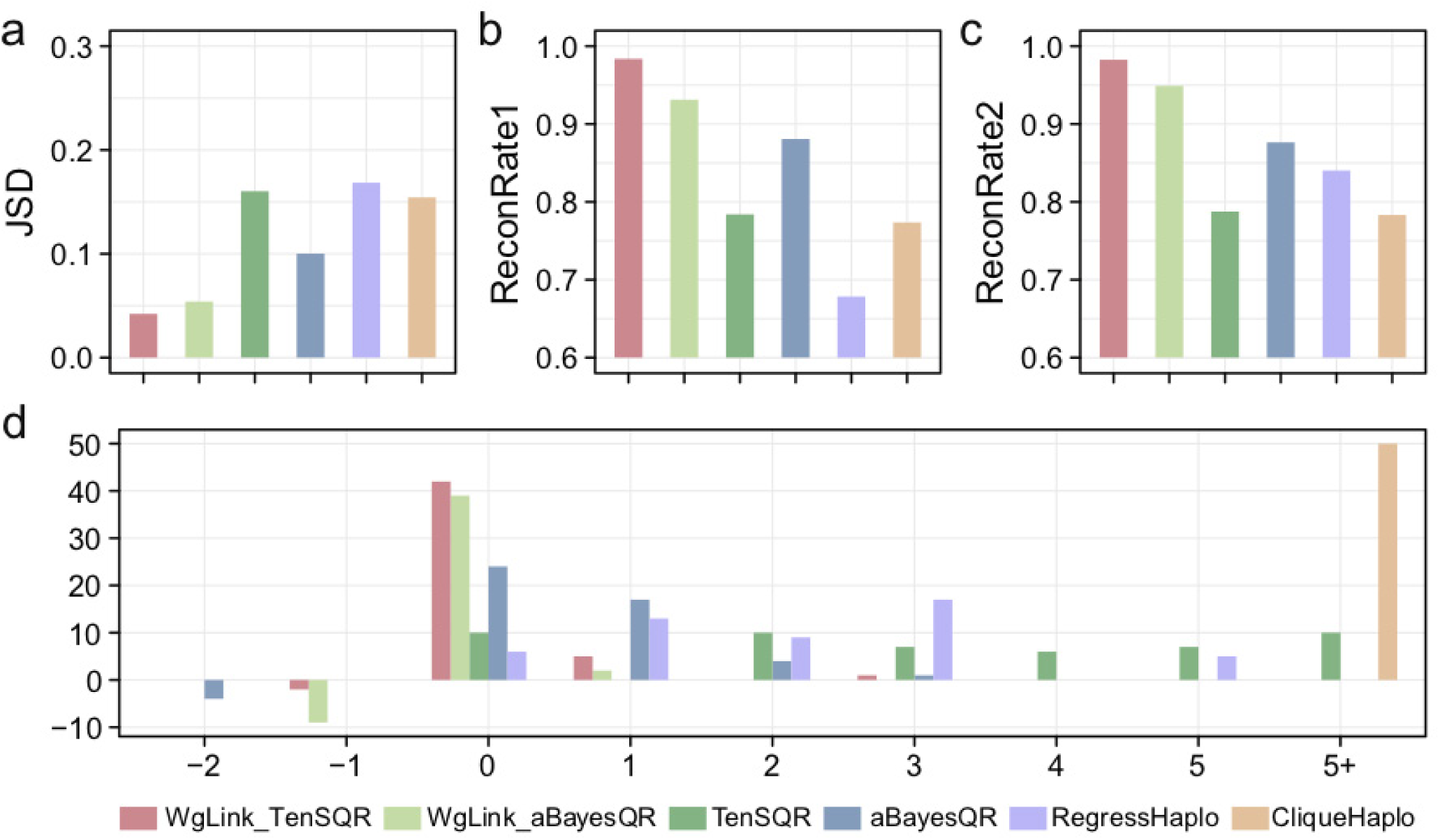
Comparison between different viral haplotype reconstruction tools based on the parameterization: 150bp read length, 10000X coverage, 0.1% sequencing error rate. Fifty rounds of simulations were conducted, and the averages were plotted. Panel **a** shows the accuracy for haplotype frequency (JSD) between truth and predicted (the smaller the better); Panels **b** and **c** show the accuracy for haplotype reconstruction, ReconRate1 and ReconRate2 based on two strategies of mapping the identities described in “Comparison criteria” (the larger the better). Penal **d** displaces the distribution of differences between the actual number of haplotypes and the predicted number. 0 means identical the predicted is identical to the truth (i.e., no error); and 5+ means the tools predicted 5 more haplotypes than truth (indicating large error).

**Supplementary Fig. 5:**
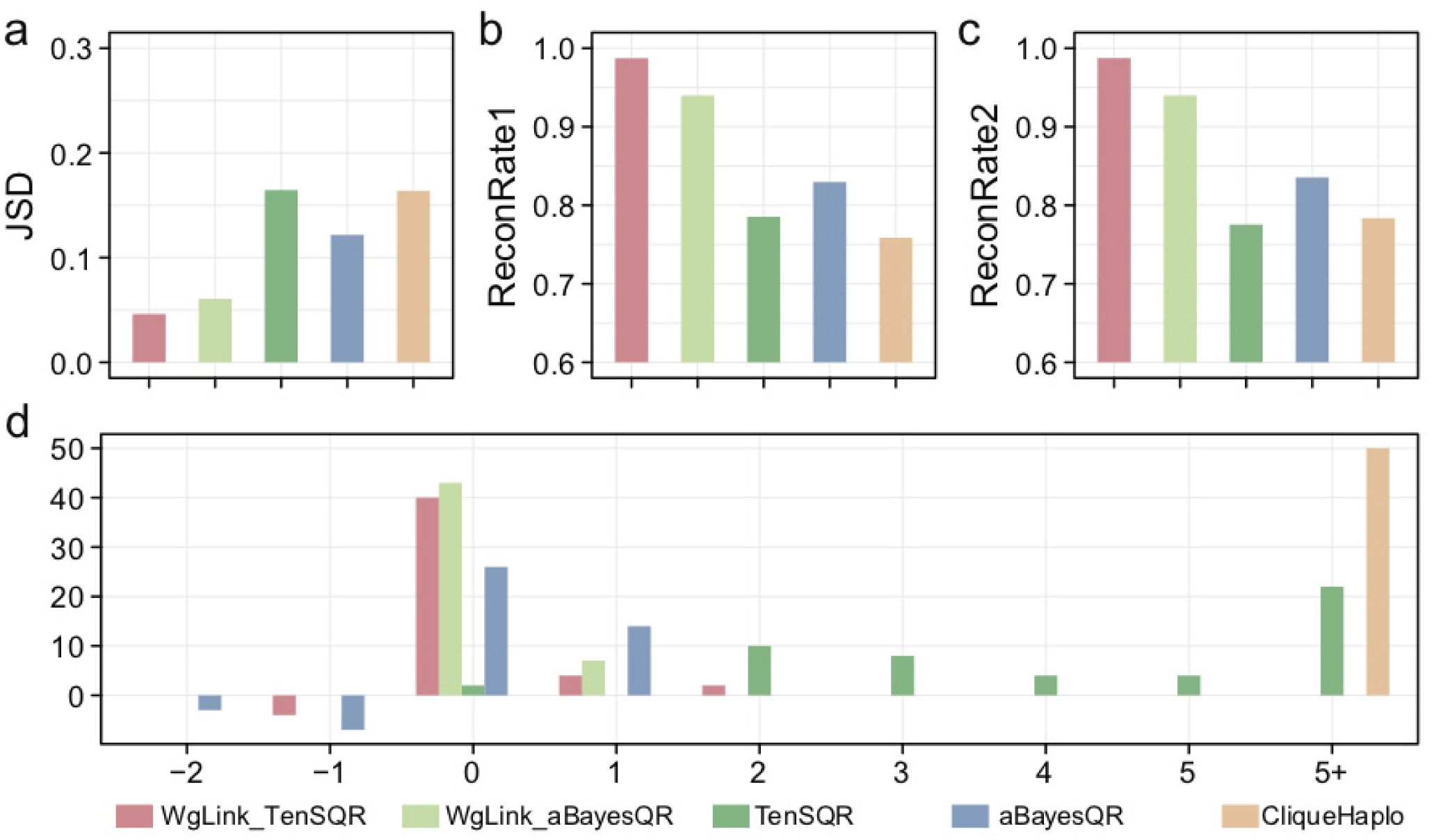
Comparison between different viral haplotype reconstruction tools based on the parameterization: 150bp read length, 10000X coverage, 0.4% sequencing error rate. Fifty rounds of simulations were conducted, and the averages were plotted. Panel **a** shows the accuracy for haplotype frequency (JSD) between truth and predicted (the smaller the better); Panels **b** and **c** show the accuracy for haplotype reconstruction, ReconRate1 and ReconRate2 based on two strategies of mapping the identities described in “Comparison criteria” (the larger the better). Penal **d** displaces the distribution of differences between the actual number of haplotypes and the predicted number. 0 means identical the predicted is identical to the truth (i.e., no error); and 5+ means the tools predicted 5 more haplotypes than truth (indicating large error). Only three alternative tools are displaced because RegressHaplo didn’t generate outcome when sequencing error rats is 0.4%.

**Supplementary Fig. 6:**
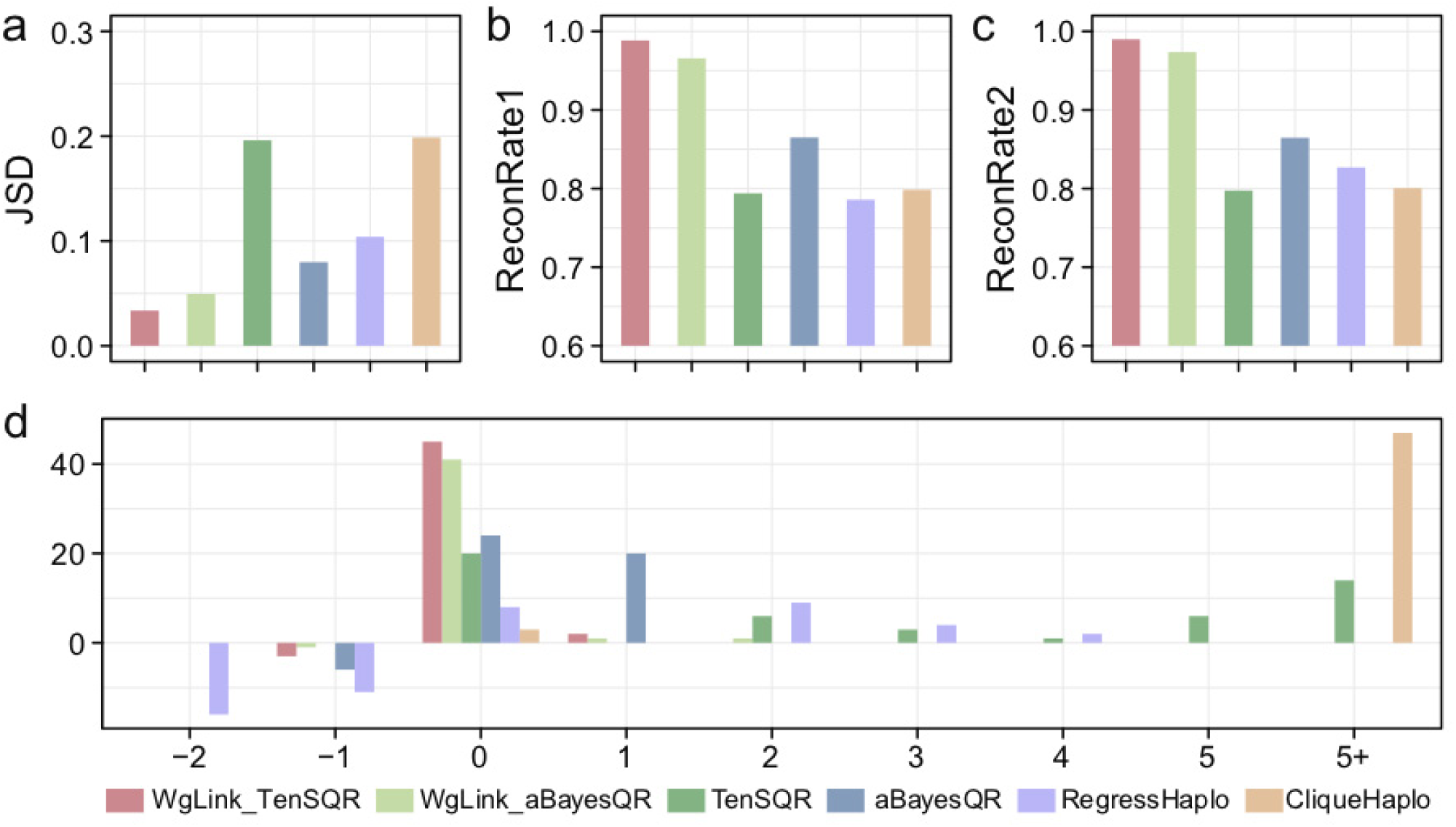
Comparison between different viral haplotype reconstruction tools based on the parameterization: 250bp read length, 2500X coverage, 0.1% sequencing error rate. Fifty rounds of simulations were conducted, and the averages were plotted. Panel **a** shows the accuracy for haplotype frequency (JSD) between truth and predicted (the smaller the better); Panels **b** and **c** show the accuracy for haplotype reconstruction, ReconRate1 and ReconRate2 based on two strategies of mapping the identities described in “Comparison criteria” (the larger the better). Penal **d** displaces the distribution of differences between the actual number of haplotypes and the predicted number. 0 means identical the predicted is identical to the truth (i.e., no error); and 5+ means the tools predicted 5 more haplotypes than truth (indicating large error).

**Supplementary Fig. 7:**
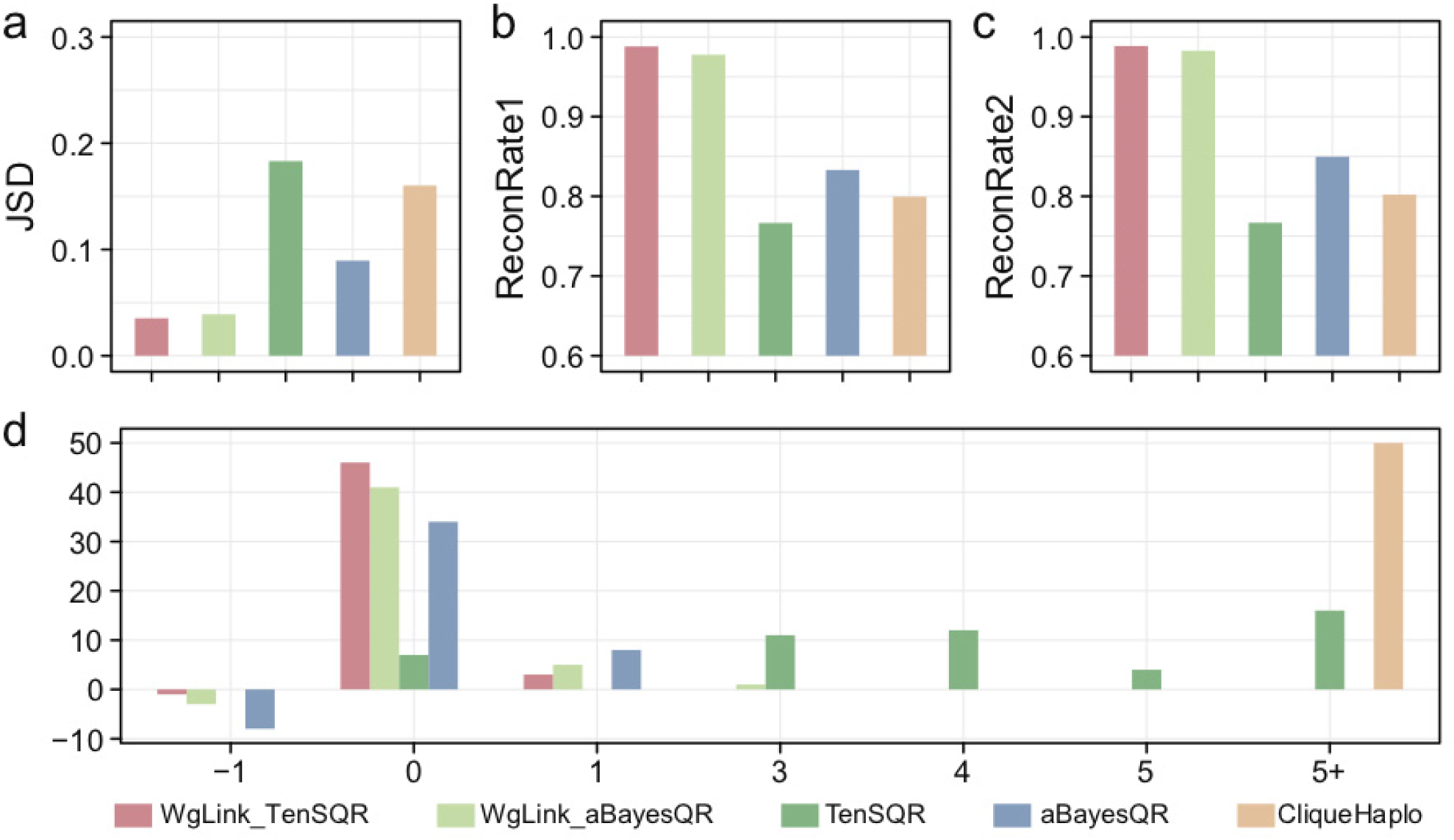
Comparison between different viral haplotype reconstruction tools based on the parameterization: 250bp read length, 2500X coverage, 0.4% sequencing error rate. Fifty rounds of simulations were conducted, and the averages were plotted. Panel **a** shows the accuracy for haplotype frequency (JSD) between truth and predicted (the smaller the better); Panels **b** and **c** show the accuracy for haplotype reconstruction, ReconRate1 and ReconRate2 based on two strategies of mapping the identities described in “Comparison criteria” (the larger the better). Penal **d** displaces the distribution of differences between the actual number of haplotypes and the predicted number. 0 means identical the predicted is identical to the truth (i.e., no error); and 5+ means the tools predicted 5 more haplotypes than truth (indicating large error). Only three alternative tools are displaced because RegressHaplo didn’t generate outcome when sequencing error rats is 0.4%.

**Supplementary Fig. 8:**
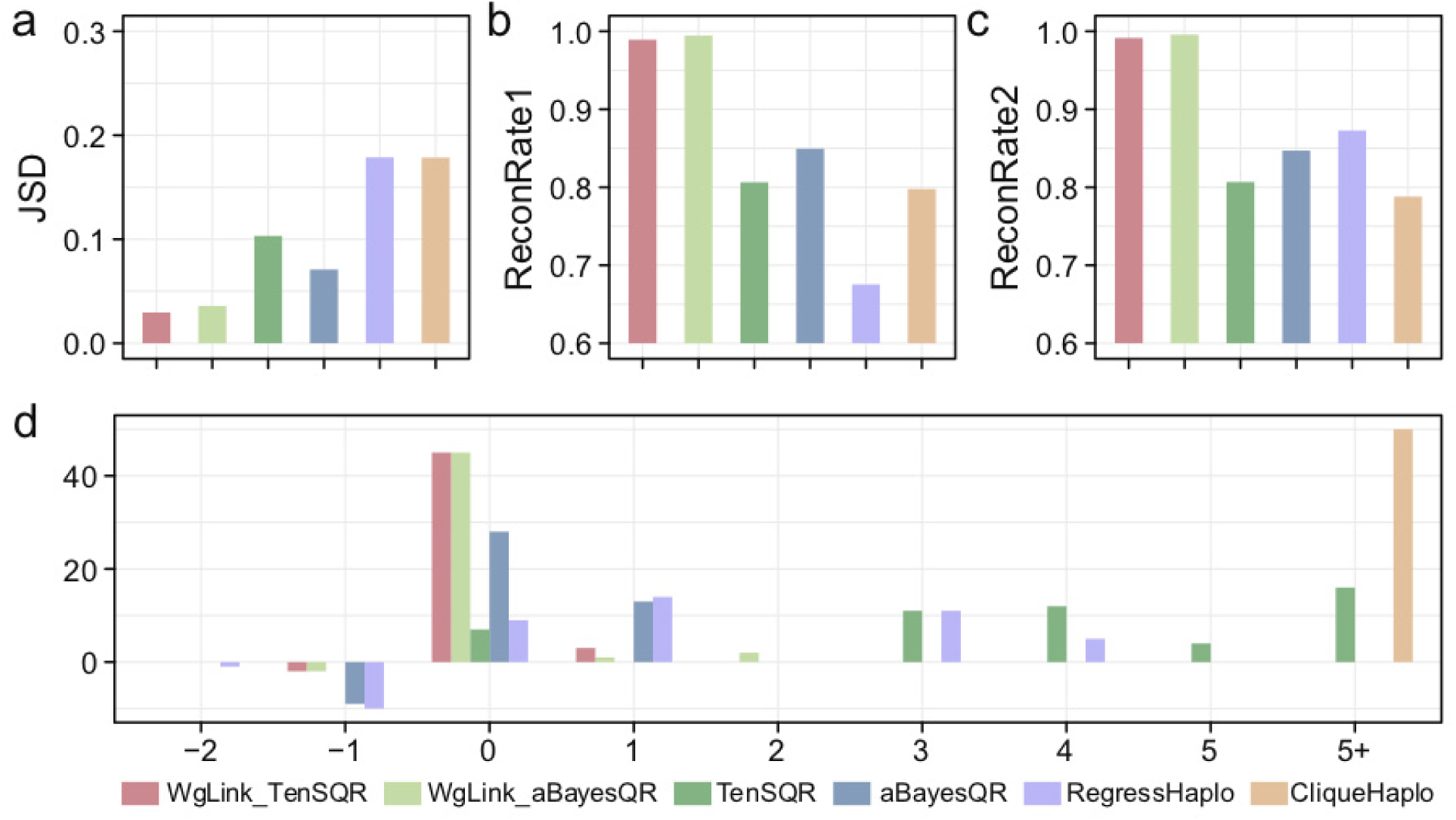
Comparison between different viral haplotype reconstruction tools based on the parameterization: 250bp read length, 10000X coverage, 0.1% sequencing error rate. Fifty rounds of simulations were conducted, and the averages were plotted. Panel **a** shows the accuracy for haplotype frequency (JSD) between truth and predicted (the smaller the better); Panels **b** and **c** show the accuracy for haplotype reconstruction, ReconRate1 and ReconRate2 based on two strategies of mapping the identities described in “Comparison criteria” (the larger the better). Penal **d** displaces the distribution of differences between the actual number of haplotypes and the predicted number. 0 means identical the predicted is identical to the truth (i.e., no error); and 5+ means the tools predicted 5 more haplotypes than truth (indicating large error).

**Supplementary Fig. 9:**
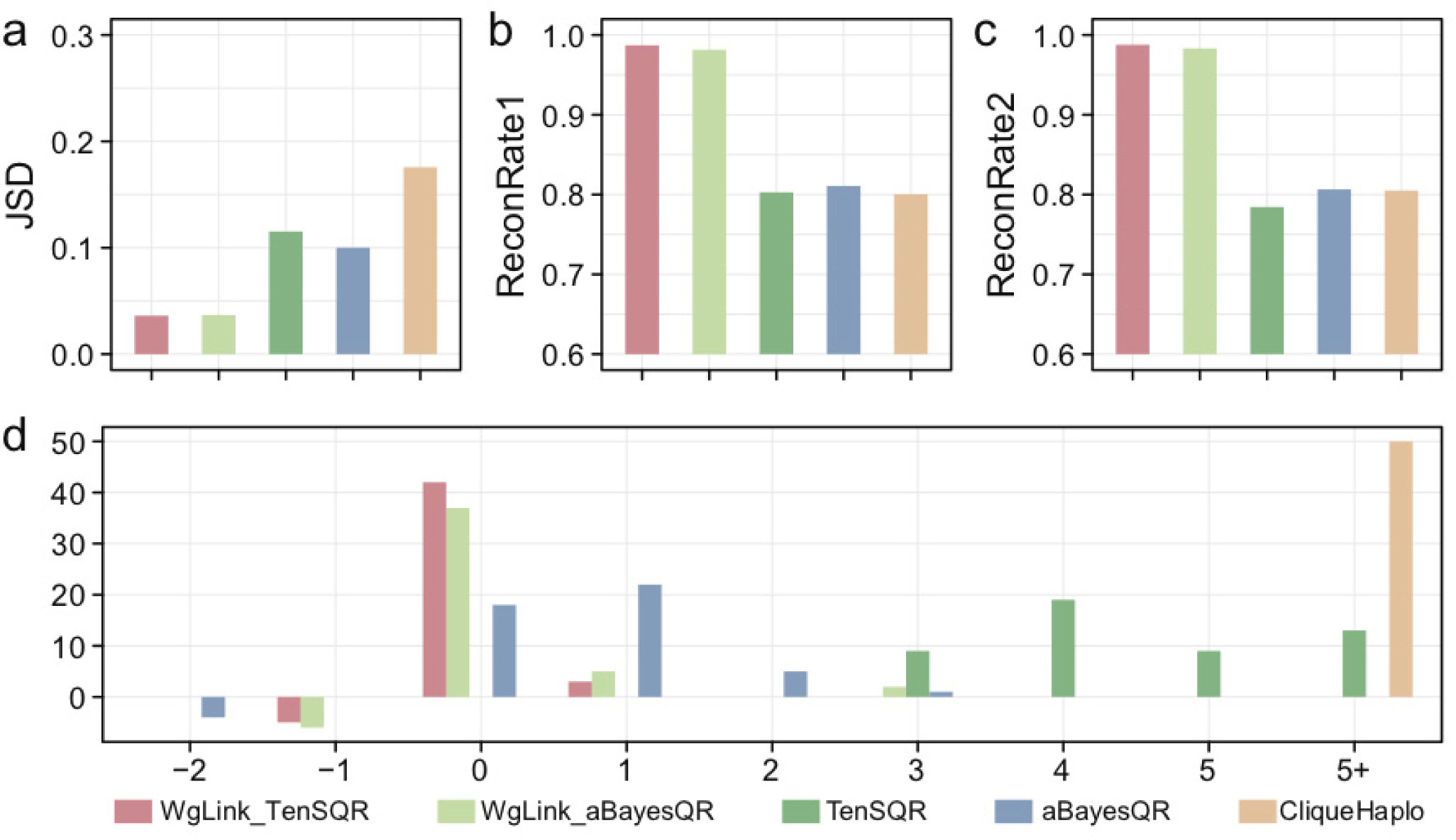
Comparison between different viral haplotype reconstruction tools based on the parameterization: 250bp read length, 10000X coverage, 0.4% sequencing error rate. Fifty rounds of simulations were conducted, and the averages were plotted. Panel **a** shows the accuracy for haplotype frequency (JSD) between truth and predicted (the smaller the better); Panels **b** and **c** show the accuracy for haplotype reconstruction, ReconRate1 and ReconRate2 based on two strategies of mapping the identities described in “Comparison criteria” (the larger the better). Penal **d** displaces the distribution of differences between the actual number of haplotypes and the predicted number. 0 means identical the predicted is identical to the truth (i.e., no error); and 5+ means the tools predicted 5 more haplotypes than truth (indicating large error). Only three alternative tools are displaced because RegressHaplo didn’t generate outcome when sequencing error rats is 0.4%.

**Supplementary Table 1:** Memory usage and computation time of different reconstruction tools. The memory usage and computation time are calculated using the simulations data for read length = 150bp, sequencing coverage = 2500X, and sequencing error rate = 0.1%. Among the usage since 50 rounds of simulations, we chose to present the average computation time and maximum memory. Multiple threads (8 threads) are used by WgLink_TenSQR, WgLink_aBayesQR, and CliqueSNV. To our knowledge, the other tools do not support multithreading.

| Tools | Time(min) | Maximum Memory (GB) |
| --- | --- | --- |
| WgLink_TenSQR | 23+5 | 3 |
| WgLink_aBayesQR | 19+5 | 8 |
| TenSQR | 79 | 9 |
| aBayesQR | 1394 | 5 |
| RegressHaplo | 157 | 76 |
| CliqueSNV | 245 | 54 |

**Supplementary Table 2:** Comparison of different reconstruction tools on the real 5 strain mixture dataset. RegreeHaplo did not successfully output the result and the computation time of aBayesQR exceeded the maximum allowed time in our HPC server. (7 days). The haplotype frequency for the 5 strains.

| Tools | Number of Predicted Haplotypes | ReconRate1(%) | ReconRate2(%) | JSD |
| --- | --- | --- | --- | --- |
| WgLink_TenSQR | 5 | 97.2 | 97.2 | 0.04 |
| WgLink_aBayesQR | 5 | 93.7 | 93.7 | 0.08 |
| TenSQR | 10 | 73.7 | 76.1 | 0.29 |
| CliqueSNV | 12 | 80.9 | 80.2 | 0.18 |
| **<u>Perfect estimates</u>** | **<u>5</u>** | **<u>100</u>** | **<u>100</u>** | **<u>0</u>** |

